# Wide-field optical redox imaging with leading-edge detection for assessment of patient-derived cancer organoids

**DOI:** 10.1101/2024.12.23.630148

**Authors:** Amani Gillette, Shirsa Udgata, Alexa E. Schmitz, Jordan E. Stoecker, Dustin Deming, Melissa Skala

## Abstract

Patient-derived cancer organoids (PDCOs) are a valuable model to recapitulate human disease in culture with important implications for drug development. However, current methods for assessing PDCOs are limited. Label-free imaging methods are a promising tool to measure organoid level heterogeneity and rapidly screen drug response in PDCOs. The aim of this study was to assess and predict PDCO response to treatments based on mutational profiles using label-free wide-field optical redox imaging (WF ORI). WF ORI provides organoid-level measurements of treatment response without labels or additional reagents by measuring the autofluorescence intensity of the metabolic co-enzymes NAD(P)H and FAD. The optical redox ratio is defined as the fluorescence intensity of [NAD(P)H / NAD(P)H +FAD] which measures the oxidation-reduction state of PDCOs. We have implemented WF ORI and developed novel leading-edge analysis tools to maximize the sensitivity and reproducibility of treatment response measurements in colorectal PDCOs. Leading-edge analysis improves sensitivity to redox changes in treated PDCOs (GΔ = 1.462 vs GΔ = 1.233). Additionally, WF ORI resolves FOLFOX treatment effects across all PDCOs better than two-photon ORI, with ∼7X increase in effect size (GΔ = 1.462 vs GΔ = 0.189). WF ORI distinguishes metabolic differences based on driver mutations in CRC PDCOs identifying *KRAS+PIK3CA* double mutant PDCOs vs wildtype PDCOs with 80% accuracy and can identify treatment resistant mutations in mixed PDCO cultures (GΔ = 1.39). Overall, WF ORI enables rapid, sensitive, and reproducible measurements of treatment response and heterogeneity in colorectal PDCOs that will impact patient management, clinical trials, and preclinical drug development.

**Statement of Significance:** Label-free wide-field optical redox imaging of patient-derived cancer organoids enables rapid, sensitive, and reproducible measurements of treatment response and heterogeneity that will impact patient management, clinical trials, and preclinical drug development.

## 1. Introduction

Precision medicine seeks to enhance cancer treatment by aligning the best therapies to individual patients, often relying on genomic mutations for decision-making^1–3^. While this approach works well for certain mutations, like *EGFR* in lung cancer and *BRAF* in melanoma, it has limitations across other cancers^4^. Patient-derived cancer organoids (PDCOs) are three-dimensional cultures created from fresh tumor specimens and retain key biologic features of the individual patient’s cancer from which they were derived. This method provides functional insights into drug sensitivity, complementing genomic analyses. PDCOs effectively replicate the molecular, histopathological, and phenotypic characteristics of the original tumors, offering accurate models for assessing patient drug responses^5–7^. They also reflect the cellular diversity within tumors, which is crucial for understanding treatment failures related to resistant cell populations^8–10^. In colorectal cancer (CRC), there are multiple molecularly defined subtypes with targeted therapy options and resistance can develop through the acquisition or selection for clones with resistant molecular profiles^6,10–12^.Consequently, evaluating drug responses across multiple PDCOs is essential for capturing treatment response within a patient. ^6^. However, traditional assessment methods used for two-dimensional cultures do not translate well to the three-dimensional nature of PDCOs. Most available PDCO assays measure metabolic activity of the cells at the well-level, limiting the ability to evaluate the heterogeneity of individual PDCOs, and therefore, heterogeneity within a patient^13–16^. Additionally, longitudinal monitoring is often difficult as these assays are highly sensitive to baseline plating of the organoids, making it more challenging to control than with two-dimensional cultures^17–21^. These challenges emphasize the need for non-destructive methods to quantify drug responses in PDCOs while accounting for cellular heterogeneity.

Cancer alters cellular metabolism, and many therapies aim to disrupt the altered metabolism to inhibit growth or trigger cell death^22^. The autofluorescent properties of the metabolic co-enzymes NADH, NADPH, and FAD offer a non-invasive way to monitor cellular metabolism in intact samples^23–30^. NADH and NADPH have overlapping fluorescent properties and are jointly referred to as NAD(P)H. It is important to note that they do play different roles in metabolic pathways, and while the NADH/NAD+ ratio is generally less oxidized in cells compared to the NADPH/NADP+ ratio, the ratios are coupled. This suggests that changes in the combined NAD(P)H autofluorescence can be used to monitor for changes in the reduced state of the individual cofactors^31–33^. The fluorescence properties of FAD, an electron acceptor, and NAD(P)H, electron donors, are often combined into the optical redox ratio (ORR) which reflects the oxidation-reduction state of a sample. The ORR is sensitive to early metabolic changes induced by drugs, which can occur before noticeable shifts in PDCO size, cell proliferation and cell death^34–37^. However, most studies of autofluorescence in PDCOs have used expensive two-photon (2P) and/or fluorescence lifetime imaging microscopes that are not widely available^9,34,38,39^.

Label-free optical redox imaging (ORI) can also be performed using commonly available wide-field microscopes equipped with standard components such as a broadband excitation source, scientific monochrome camera, and standard 4′,6-diamidino-2-phenylindole (DAPI) and green fluorescent protein (GFP) filter cubes to collect NAD(P)H and FAD respectively^40,41^. Our group and others have previously used wide-field ORI (WF ORI) at both cellular and tissue levels to evaluate oxidative stress, mitochondrial function, and drug responses^27,42,43^. While WF ORI cannot resolve individual cells as effectively as 2P microscopy, due to the low signal of the autofluorescent co-enzymes, it allows for the rapid imaging of large PDCO populations, which is crucial for studying subpopulations with varying treatment responses ^10,44,45^. Our prior work demonstrated that WF ORI is a practical tool for monitoring morphological and metabolic changes in PDCOs, with single-organoid tracking enhancing sensitivity to drug responses^41^. We found that the autofluorescence signal at the edge of the PDCOs was different from that of the core due to differing biological environments such as the necrotic core^46^ that generally has high FAD signal^47,48^. However, our prior work did not isolate signal to the living cells on the edge of the PDCOs.

Here, we expand upon prior work showing that an additional analysis step, leading-edge detection, improves sensitivity to treatment responses. Additionally, we performed a direct comparison between matched PDCOs imaged with both WF ORI and 2P ORI. Finally, we assessed the sensitivity of WF ORI to mutation profiles and targeted therapies in a blinded study of mixed populations of PDCOs. This work demonstrates that wide-field optical redox imaging is an accessible tool for monitoring changes in metabolism in PDCOs, capable of reproducibly assessing treatment response and heterogeneity in clinically relevant treatment studies.

## 2. Methods

### 2.1 CRC PDCO isolation and organotypic culture techniques

All studies were completed following Institutional Review Board (IRB) approval with informed consent obtained from subjects through the University of Wisconsin (UW) Molecular Tumor Board Registry (UW IRB#2014-1370) or UW Translational Science BioCore (UW IRB#2016-0934). All methods were performed in accordance with the relevant guidelines and regulations. CRC tissue was obtained from needle, endoscopic biopsies or primary resections and processed as previously described^49,50^. After processing, PDCO suspensions were immediately mixed at a 1:1 ratio with Matrigel (Corning) and conditioned media. Droplet suspensions were plated and set for three to five minutes at 37°C then inverted for at least 30 minutes to solidify the matrix and avoid the direct interface of PDCOs with the interface. Plated cultures were overlaid with 450μL of media, DMEM/F12 (Invitrogen) with added 1x Glutamax (Invitrogen), 10mM HEPES (Fisher), 50IU/mL penicillin-streptomycin (Invitrogen), 50ng/mL EGF (Invitrogen), mixed at a 1:1 ratio with WNT3a conditioned media^49^. PDCOs were incubated at 37°C in 5% CO2 and medium was replaced every 48-72 hours.

### 2.2 Therapeutic Studies

PDCOs were collected from 24-well culture plates and passaged to single well gridded 35mm glass dishes (Cellvis) or 24-well glass plates. Images were taken on a Nikon Ti-2E inverted microscope using a 4x objective prior to treatment as described below. Pharmacologic agents were prepared at physiologic C_max_ and the duration of therapy was 48 hours for FOLFOX studies (10μM 5-FU + 5μM Oxaliplatin)^51,52^ followed by 48 hours in media prior to imaging, whereas and panitumumab^53^ (1.6μM) PDCOs were treated for 48 hours and imaged at 48 hours. Pharmacologic agents were received from the University of Wisconsin Carbone Cancer Center Pharmacy including 5-fluorouracil (5-FU, Fresenius), oxaliplatin (Hospira), and panitumumab (Amgen). All therapeutic studies were performed with two to three independent biological replicates.

### 2.3 CRC mutation specific PDCO Mixing Studies and Predictions

CRC PDCOs with known mutation profiles, wild-type (WT) or *PIK3CA / KRAS* double mutant (DM) PDCOs, were cultured separately and then passaged via spiking, using a pipette tip to isolate a single PDCO **(Supplementary Video 1)**, and plated into mixed population Matrigel droplets. The scientist culturing immediately collected and annotated Brightfield (BF) images with the known mutation profile of the individual PDCOs in the image, prior to collecting WF ORI for day 0. The image analysis scientist was not given the BF images, and therefore was blinded to mutation profile, until after completing the image analysis and predictions.

### 2.4 WF ORI and Leading-edge Image Analysis

Widefield ORI was performed using a Nikon Ti-2E coupled to a SOLA light engine (300 to 650 nm, Lumencor, Beaverton, Oregon, United States). All images were collected with NIS Elements software (Nikon) using a 4x air, 0.13 NA objective (Plan Apo, Nikon), and Hamamatsu Flash4 digital CMOS camera. NAD(P)H was excited for 3s through a 360/40 nm filter at 40% power (0.25 mJ/cm^2^), and emissions were collected using a 400 nm dichroic mirror and a standard 4′,6-diamidino-2-phenylindole (DAPI, 460/50 nm) filter. FAD was sequentially excited for 3.5s through a 480/30 nm filter at 40% power (0.531 mJ/cm^2^) and emissions collected using a 505 nm dichroic mirror and a standard fluorescein isothiocyanate (FITC, 535/20 nm) filter. Five FOVs (image size = 3.34mm x 3.34 mm; 2044 x 2048 pixels) were taken of each sample, with three replicates per sample condition, light doses remain below levels shown to keep PDCOs viable ^40,54^. Images were acquired on Day 0 prior to treatment and post treatment (Day 4 for FOLFOX studies, Day 2 for Panitumumab studies) for all PDCOs, using the same settings.

NAD(P)H and FAD widefield autofluorescence images were analyzed using CellProfiler^55^. Due to the biological heterogeneity within PDCO’s which often contain necrotic cores, quiescent zones, and proliferating zones **(Fig. 1A)**, especially in large PDCOs, a pipeline to limit ORI measurements to the proliferating zone of the PDCOs called “leading-edge analysis” was developed **(Fig. 1B)**. PDCOs were segmented by hand using the NAD(P)H channel. The radius of the individual PDCO masks was then calculated by CellProfiler, and 20 pixels (60 μm) were subtracted generating a core mask, which was subtracted from the original PDCO mask to create a leading-edge mask **(Fig. 1B)**. The optical redox ratio (ORR) was then calculated for each leading-edge mask, we expect day-to-day variation in the widefield system (e.g., minor drift, variation in exposure time), and recognize that illumination through the slightly autofluorescent Matrigel may affect intensity-based measurements. To account for these factors, we measured the average fluorescence intensity from five background (PDCO-free) locations within each image. We calculated a background-normalized redox ratio for all PDCOs on each image as follows:

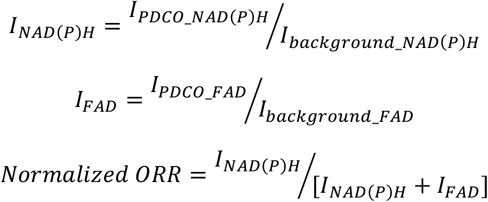

**Figure 1.**
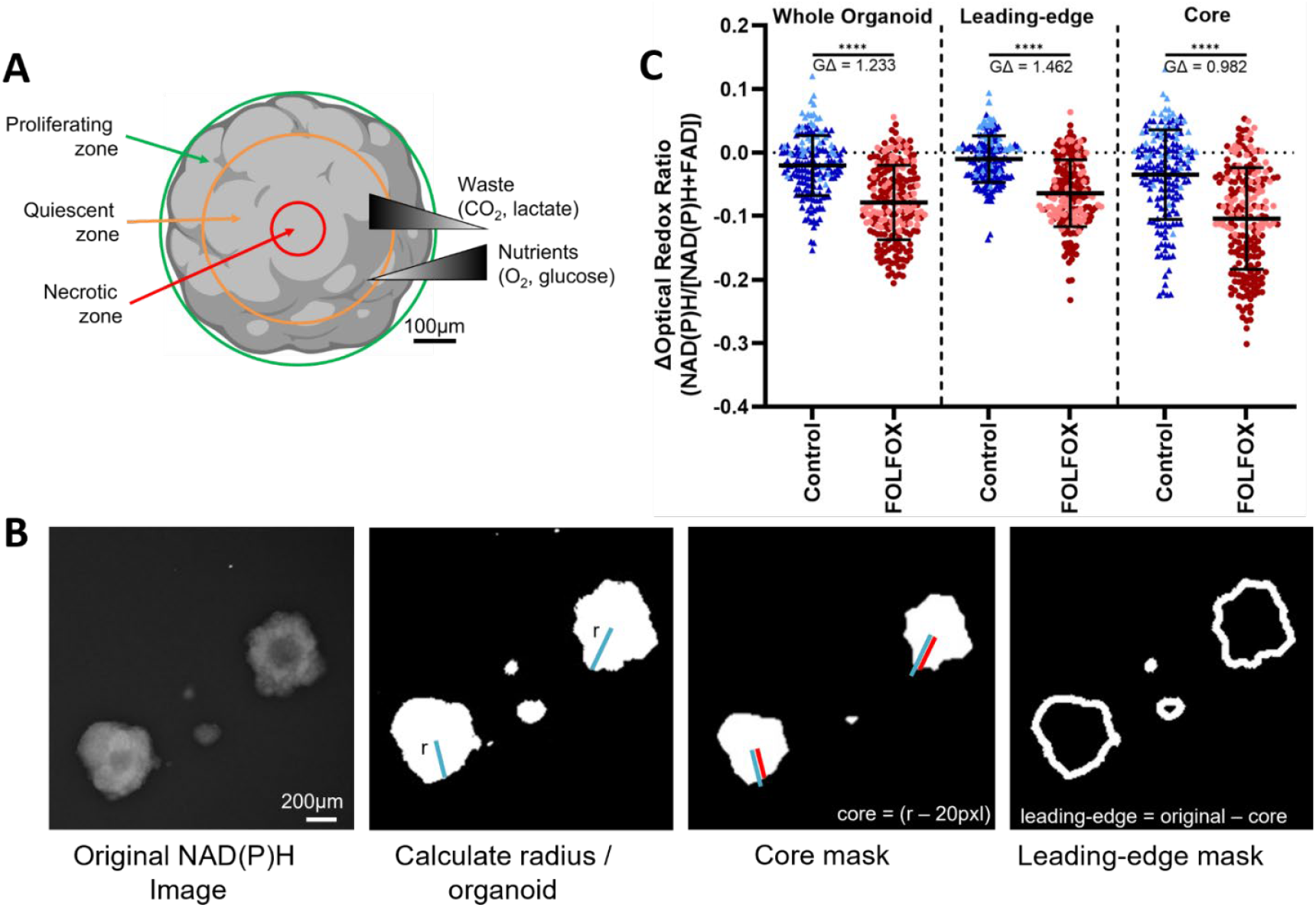
Leading-edge analysis captures redox change in proliferating zone of PDCOs. (**A**) PDCOs have different regions of proliferating (green), quiescent (orange), and dead (red) cells, due to nutrient and waste gradients. (**B**) Leading-edge analysis isolates redox changes within the proliferating zone by calculating a PDCO mask from the original wide-field NAD(P)H intensity image, defining the radius (blue line) of each PDCO, then 20 pixels (60μm) are subtracted from the radius to generate a leading-edge mask. (**C**) Change in optical redox ratio from day 0 to 4 for individual PDCOs. Leading-edge analysis focuses on cells in the proliferating zone alone, improving the sensitivity to FOLFOX (10μM 5-FU + 5μM Oxaliplatin) response compared to whole PDCO analysis. ****p<0.0001, ordinary one-way ANOVA; GΔ, effect size.

Note that other definitions of the optical redox ratio have been used in other prior papers^30,56^, and we choose this definition to be consistent with our prior work and so that the ORR decreases with effective treatments^37,49^. Treatment effects were measured by the change in the optical redox ratio (ΔORR), where ΔORR = Normalized ORR_organoid_day4_ – Normalized ORR_average_day0_. Here, “Normalized ORR_organoid_day4_” is the background normalized optical redox ratio of individual PDCO on day 4, and “Normalized ORR_average_day0_” is the background normalized average optical redox ratio of the same PDCO population measured on day 0 prior to treatment. For each condition, a minimum of 21 PDCOs were assessed with WF ORI, see **supplementary table 1** for PDCO counts per condition and experiment. Unless otherwise noted, all ΔORR values are reported using the leading-edge technique.

### 2.5 Two-Photon Imaging and Analysis

Briefly, NAD(P)H and FAD were excited at 750 nm and 890 nm, respectively, using a tunable ultrafast pulsed laser (Coherent, Inc), an inverted microscope (Nikon, Eclipse Ti), and a 40x water immersion (1.15NA, Nikon) objective. Both NAD(P)H and FAD images were obtained for the same field of view. FAD fluorescence was isolated using an emission bandpass filter of 550/100 nm, while NAD(P)H fluorescence was isolated using an emission bandpass filter of 440/80 nm. Fluorescence lifetime images were collected using time-correlated single photon counting electronics (SPC-150, Becker and Hickl) and a GaAsP photomultiplier tube (H7422P-40, Hamamatsu). Images (512 x 512 pixels) were obtained using a pixel dwell time of 4.8μs over 60s total integration time. A Fluoresbrite YG microsphere (Polysciences Inc.) was imaged as a daily standard for fluorescence lifetime. The instrument response function was measured using a second-harmonic generation signal from urea crystals excited at 890 nm, with full width at half-maximum of 240 ps. The lifetime decay curves were fit to a single exponential decay, and the fluorescence lifetime was measured to be 2.1 ns (*n* = 7), which is consistent with published values^9,37^.

Fluorescence intensity was computed for each image pixel using SPCImage (v8.0, Becker & Hickl GmbH). To enhance the fluorescence counts in each decay, a 3×3 bin comprising nine neighboring pixels was employed (>1000 photons/pixel) ^57^. The background was thresholded, and the pixel-wise decay curves were fit to a biexponential model convolved with the instrument response function, using an iterative parameter optimization to obtain the lowest sum of the squared differences between the model and data (weighted least squares algorithm). The decay curves were then integrated to generate the intensity of each fluorophore. Here, we only report the intensity values, and the lifetime parameters are not reported.

Whole-cell masks were generated using CellProfiler as previously described^58^. Briefly, pixels belonging to nuclear regions were manually identified and the resulting round objects were stored as a mask. Cells were identified by propagating out from each nucleus, an Otsu Global threshold was used to improve the propagation and prevent it from continuing into background pixels. Cell cytoplasm was defined as the cell border minus the nuclei. Values for intensity of NAD(P)H and FAD as well as the ORR were measured for each cell cytoplasm. For each variable, values for all cells within a treatment group were pooled together, 426 to 1548 cells were analyzed per condition, see **supplementary table 2** for cell counts per condition.

### 2.6 Statistical Analysis

A post-hoc power analysis of the ΔORR for control vs FOLFOX treated PDCOs was performed across a strong and weak responding PDCO line to determine sample sizes needed for continued experiments, using an alpha of 0.01, a statistical power of 0.95, and Glass’s Delta effect size measurements, described below (**Table 1**).

**Table 1.** Post-hoc power analysis of ΔORR for control vs FOLFOX treated PDCOs from two separate responsive lines, one with a weak response and one with a strong response.

| Sample | Control (ΔORR) | | FOLFOX (ΔORR) | | Power Analysis | | | N needed |
| --- | --- | --- | --- | --- | --- | --- | --- | --- |
|  | Mean | SD | Mean | SD | alpha | Power | Effect Size (GD) |  |
| Strong Response PDCO line | 0.0191 | 0.0339 | -0.0661 | 0.0426 | 0.01 | 0.95 | 2.53 | 8 |
| Weak Response PDCO line | -0.0196 | 0.0326 | -0.0592 | 0.0533 | 0.01 | 0.95 | 1.33 | 25 |

Differences in ΔORR between treatment and control organoids were tested using a one-way ANOVA of control vs. treatment, with a p-value < 0.05 defined as significant. Treatment effect size was calculated using Glass’s Delta (GΔ) which is defined as:

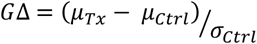

where μ_Tx_ is the mean ΔORR of the treatment group, μ_Ctrl_ is the mean ΔORR of the control group, and σ_Ctrl_ is the standard deviation of the ΔORR for the control group^59,60^. GΔ was used because comparisons of large sample sizes, such as those acquired here with single cell and single PDCO measurements, almost always pass traditional significance tests unless the population effect size is near zero.

## 3. Results

### 3.1 Leading-edge analysis improves sensitivity to redox change in treated PDCOs

PDCOs are known to have different biological zones that correspond to differing cell states primarily driven by nutrient access. The diffusion limit of most small molecules, such as oxygen and ATP, is around 100-150μm into the organoid, which is similar to that in a vascular tissue and tumor masses *in vivo*^61,62^. This means that the outermost edge of the PDCO with the most exposure to nutrients is dominated by proliferative cells with active metabolism. Cells that are exposed to high levels of metabolic waste (CO_2_ and lactate) and starvation due to lack of nutrient diffusion exist in the necrotic zone, while the quiescent zone between the proliferative and necrotic zones contains live cells that are not highly metabolically active **(Fig. 1A)**^62,63^. Therefore, ORR measurements can be limited to the leading-edge of the PDCO that contains only cells within the proliferating zone. This was achieved by segmenting the whole PDCO, then subtracting 20pixels (60μm) radially to generate a core mask. This core mask was then subtracted from the original PDCO mask to create a leading-edge mask of the outermost 60μm of the PDCO **(Fig. 1B)**. A 60μm threshold was chosen to conservatively remove the quiescent zone and retain the most proliferative outer edge of each PDCO.

The change in optical redox ratio (ΔORR) on day 4 post-treatment to day 0 pre-treatment for control vs FOLFOX (10μM 5-FU + 5μM Oxaliplatin) treated PDCOs is shown across two patient samples **(Fig. 1C)**. There were statistically significant changes in all measured compartments including the whole organoid, leading-edge, and core. However, the effect size of the leading-edge ΔORR was the largest (GΔ = 1.462) compared with the whole organoid and core measurements (GΔ = 1.233, GΔ = 0.982 respectively). The lower GΔ in the core mask was likely the cause of the decreased GΔ in the whole organoid mask. Overall, these results indicate some benefit to isolating the ORR to the proliferating zone of the PDCOs imaged with WF ORI.

### 3.2 WF ORI resolves treatment effects across all PDCOs better than 2P ORI

Using gridded dishes, identical PDCOs were imaged on day 0 and day 4 under control and FOLFOX treatment conditions for two patients, using both two-photon (2P) and wide-field (WF) ORI. Representative day 4 NAD(P)H autofluorescence images of control and treated PDCOs show the difference in scale and PDCO detail that can be achieved with the two imaging modalities **(Fig. 2A-B**). Boxed PDCOs in the WF panels correspond with the representative 2P images shown. Tracking the same PDCO over time with treatment using both WF and 2P ORI indicates that both modalities show an overall decrease in the ORR of the PDCOs with FOLFOX treatment **(Fig. 2C-D**), a change that has been previously correlated with treatment response^37,41,49^.

**Figure 2.**
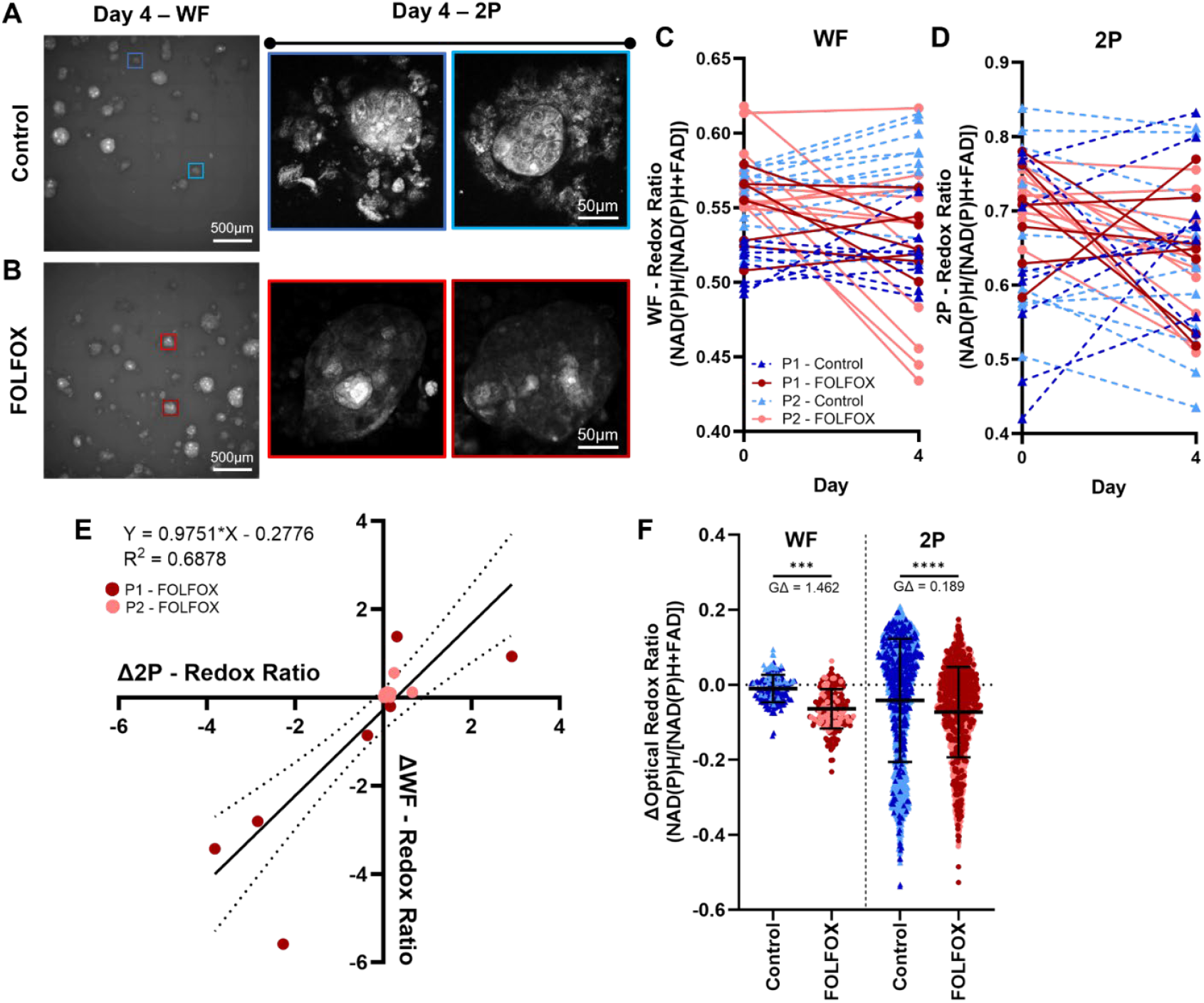
Correlation and effect sizes for two-photon (2P) and widefield (WF) optical redox imaging (ORI). Representative Day 4 images of PDCOs taken with WF and 2P NAD(P)H imaging in (**A**) Control and (**B**) FOLFOX (10μM 5-FU + 5μM Oxaliplatin) treated PDCOs, with matched 2P images of the boxed PDCOs (blue, control; red, FOLFOX). Optical redox ratio (NAD(P)H / [NADH(P)H+FAD]) decreases from Day 0 to Day 4 in matched PDCOs (n=16) imaged with both (**C**) WF and (**D**) 2P. Greater decreases were observed using WF compared to 2P. (**E**) Correlation of change in optical redox ratio from day 0 to 4 (normalized to change in controls) shows moderate correlation between 2P and WF (R^2^=0.68) in matched PDCOs (n=16). (**F**) Effect sizes differ between 2P of single cells from 16 PDCOs (1644-1974 cells / condition) and WF from all PDCOs in a dish (175-205 PDCOs /condition). ****p<0.0001, ordinary one-way ANOVA; GΔ, effect size. P1, patient 1; P2, patient 2.

By correlating the ΔORR for both 2P and WF ORI of identical PDCOs (n=16) over 4 days, (**Fig. 2E**), we see a moderate correlation (R^2^ = 0.68), which is expected given the differences in resolution and optical sectioning between WF and 2P ORI. Importantly, WF ORI can rapidly image all PDCOs in a dish (40+ PDCO / dish / 30s), providing greater power to resolve treatment effects compared to much slower 2P ORI (3-5 PDCO / dish / 30min). Specifically, WF ORI of individual PDCOs provided ∼7X increase in effect size (GΔ = 1.462) compared to single cell 2P ORI (GΔ = 0.189) for FOLFOX treatment of the same samples (**Fig. 2F**). This is significant because WF ORI can be performed with hardware present in most research and clinical labs, at lower cost and higher throughput compared to 2P ORI, with increased sensitivity to treatment.

### 3.3 WF ORI distinguishes metabolic differences based on driver mutations in CRC PDCOs

Prior work has shown that ORI is sensitive to driver mutations in breast cancer cell lines^29,64^. To investigate whether CRC PDCOs with known driver mutations exhibit differing baseline metabolic characteristics, we used WF ORI to image PDCOs across multiple mutation profiles including wild-type (WT, 4 patients), *KRAS* (3 patients), *PIK3CA* (2 patients), or both *KRAS* and *PIK3CA* (2 patients) mutants (**Fig. 3A-B**). Remarkably, these pre-treatment ORR measurements showed differences between WT and *PIK3CA* mutant PDCOs (GΔ = 0.546), and *KRAS + PIK3CA* mutant PDCOs (GΔ = 1.24). The ORR differences between WT and *KRAS + PIK3CA* double-mutant (DM) PDCOs were sufficient to classify the two populations in mixed culture (**Fig. 3C** and **Supplemental Video 1**). A random forest classifier trained on the ORR, NAD(P)H intensity, and FAD intensity from WT and DM PDCOs in pure culture and tested on dishes with a mixed population of WT and DM PDCOs achieved high sensitivity and specificity with an area under curve (AUC) of 0.87 when predicting WT vs DM PDCOs (**Fig.3D-F**). Overall, this indicates that in some cases, WF ORI can identify driver mutations pre-treatment, and could be used as a tool to screen patients for resistant cell populations especially in the case of therapies that target specific mutations.

**Figure 3.**
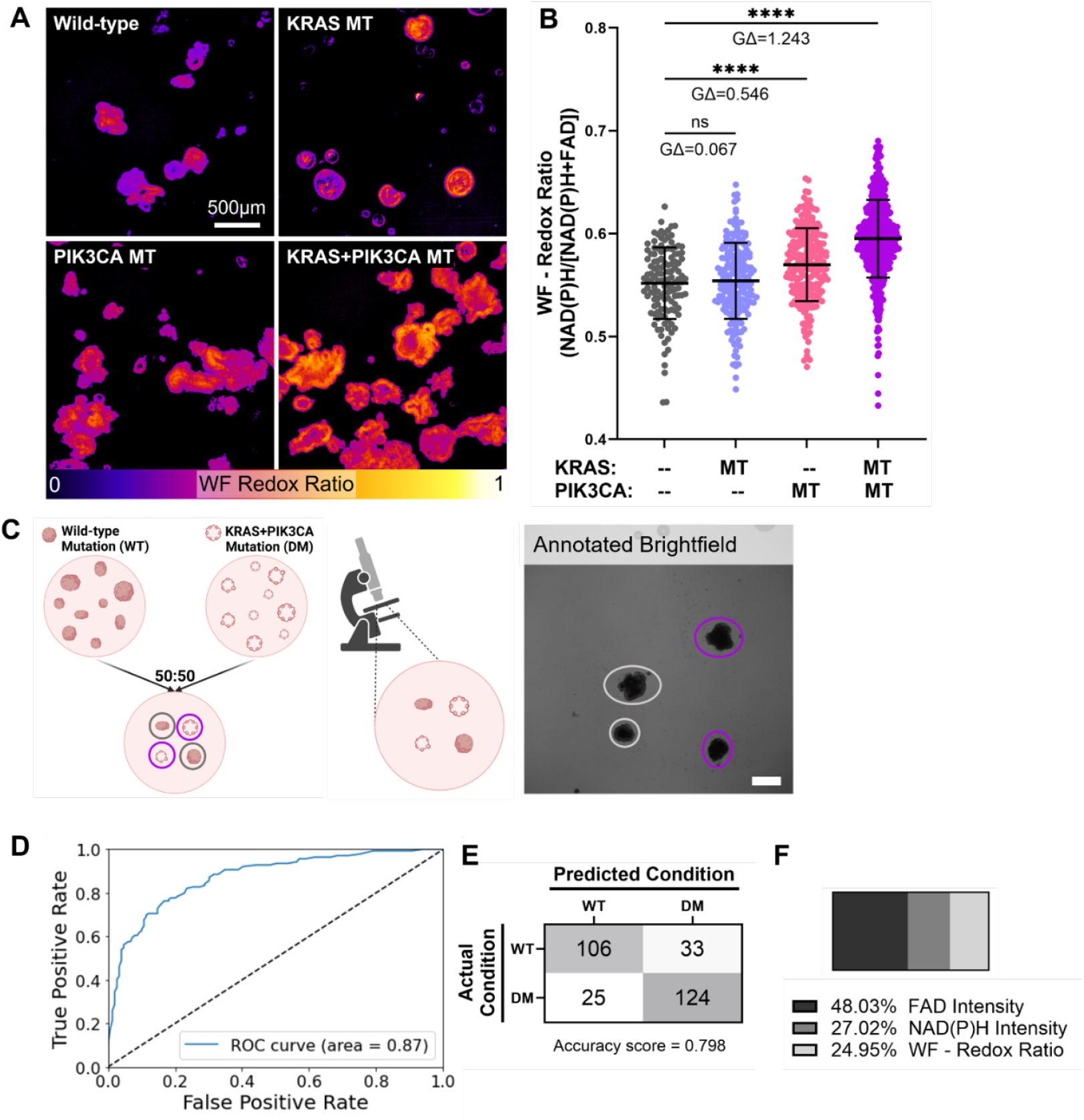
WF ORI distinguishes mutations (MT) from wild-type (WT) in CRC PDCOs. (**A**) Representative WF images of PDCOs with different KRAS and PIK3CA mutation phenotypes. (**B**) WF ORI from untreated WT (4 patients; 128 PDCOs), and MT including KRAS (3 patients; 199 PDCOs), PIK3CA (2 patients; 226 PDCOs), and KRAS+PIK3CA (2 patients; 493 PDCOs). GΔ, effect size. **\*\*\*\***p<0.0001; one-way ANOVA with Dunnett’s multiple comparisons test (**C**) Experimental set-up to generate a mixed mutation population of PDCOs for prediction and Panitumumab treatment. PDCOs from wild-type (WT) and KRAS+PIK3CA double mutant (DM) populations were spiked into a mixed population dish, the dish was then imaged and a brightfield image was annotated by the scientist who spiked the PDCO’s. (**D**) Receiver operating characteristic (ROC) curve and area under the curve (AUC) for a random forest classifier trained on 70% of the imaged WT and KRAS+PIK3CA double mutant (DM) PDCOs (pure cultures) and tested on the remaining 30% (mixed populations) yields an AUC of 0.87 using all variables (NAD(P)H intensity, FAD Intensity and WF Redox Ratio). (**E**) Confusion matrix of the random forest classifier shows performance for classification of WT and DM PDCOs. **(F)** Chart showing the relative weight of the variables included in the random forest classifier.

### 3.4 WF ORI is more sensitive to panitumumab resistant PDCOs in mixed population dishes compared to PDCO diameter

In CRC, *K/NRAS* mutations lead to clinical therapeutic resistance to EGFR monoclonal antibodies such as panitumumab^65^. Here, we test whether WF ORI can identify treatment sensitive wildtype (WT) vs treatment resistant *KRAS+PIK3CA* (DM) PDCOs in mixed dishes (**Fig. 4A**). Dishes with both WT and DM PDCOs were treated with a physiological C_max_ of panitumumab^53^ for 2-days and the same PDCOs were tracked over time with treatment using both manual PDCO diameter measurements, as described in previous publications by our group^49,50,66^, and WF ORI (**Fig. 4A**). Results show that there is an overall decrease in the diameter of the WT PDCOs with 2 days of panitumumab treatment that is not apparent in the DM PDCOs **(Fig. 4B-C**). Here, the effect size for the diameter change over 2 days of treatment for WT compared to DM PDCOs in the same dish is GΔ = 0.92. This decrease in WT PDCO diameter has been previously correlated with treatment response^49,50,66^.

**Figure 4.**
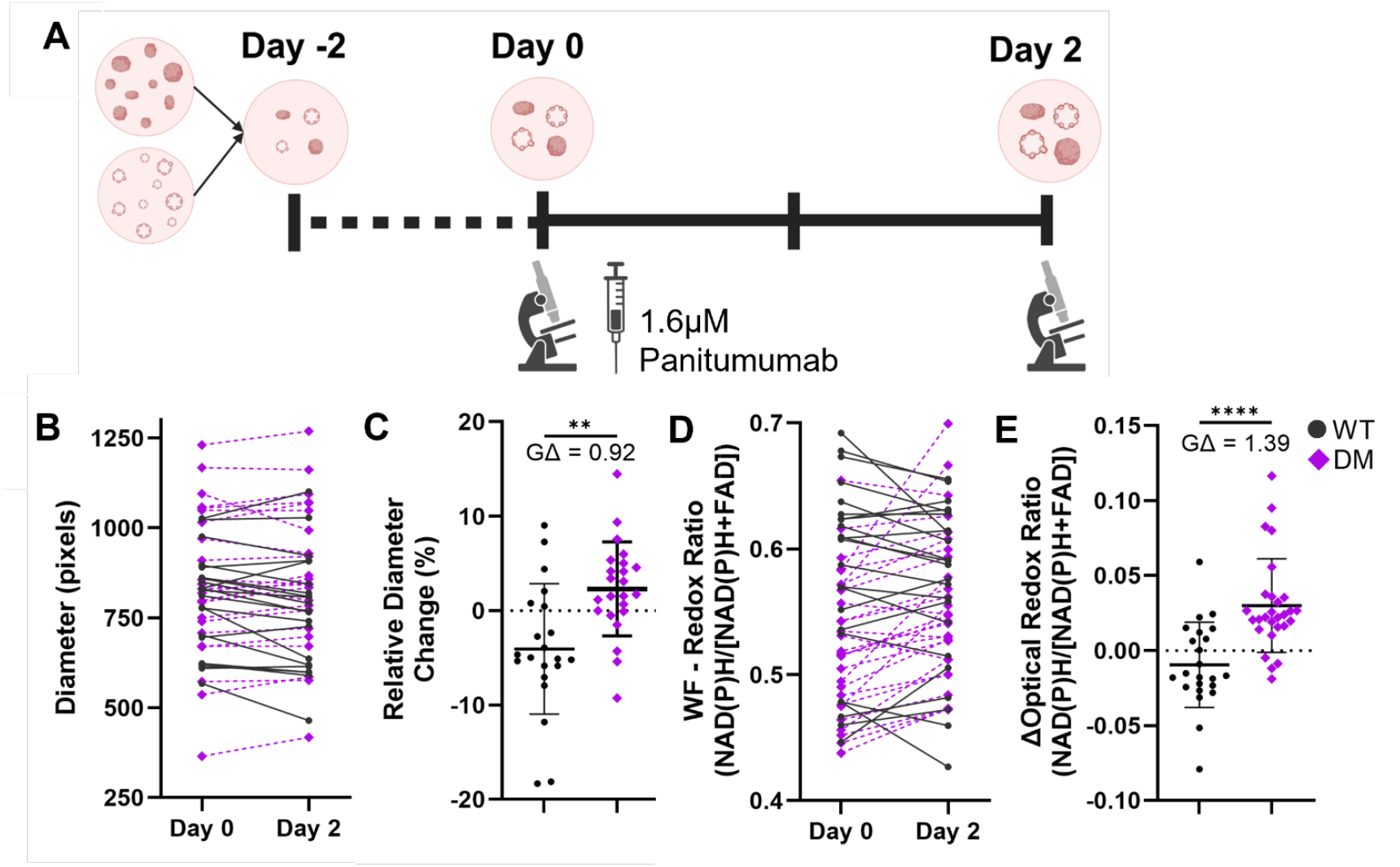
WF ORI can detect mixed mutation population of PDCOs due to targeted treatment sensitivity within 48hrs. **(A)** Experimental design, 2 days before start wild-type and KRAS+PIK3CA double mutant PDCOs were plated together, on Day 0 PDCO’s were imaged with WF and then treated with 1.6uM Panitumumab, on Day 2 PDCOs were imaged again to determine treatment response (**B**) Diameter of PDCOs from Day 0 to Day 2 in a mixed population of wild-type (WT, gray) and KRAS+PIK3CA double mutant (DM, purple) before and after panitumumab treatment. **(C)** Relative change in diameter (Day 2 – Day 0) for the individual PDCOs indicates WT PDCOs are more sensitive to panitumumab treatment than DM PDCOs **(D)** WF optical redox ratio (NAD(P)H / [NADH(P)H+FAD]) of the same PDCOs. (**E**) Change in optical redox ratio (Day 2 – Day 0) indicates that almost all WT PDCOs within a mixed dish are sensitive to panitumumab while the DM PDCOs are resistant to panitumumab. GΔ, effect size. **p<0.01, ****p<0.0001; unpaired t test with Welch’s correction.

Similarly, WF ORR shows a decrease in the ΔORR of the WT PDCOs with 2 days of panitumumab treatment that is not apparent in the DM PDCOs **(Fig. 4D-E**). A decrease in ORR has been previously correlated with treatment response^26,41,49^. However, the effect size between WT and DM PDCOs in the same dish is greater for the ΔORR (GΔ = 1.39) compared to the change in diameter (GΔ = 0.92). This demonstrates that WF ORI can rapidly assess PDCOs for mutation profiles when using therapies that target known mutational drivers of resistance. These WF ORR drug response screens could be used to improve treatment selection for individual patients.

## 4. Discussion

Functional drug screenings using patient-derived cancer organoids (PCOs) reflect the diversity and drug sensitivity of tumors *in vivo*, making them valuable tools for predicting the most effective treatments and identifying new drug candidates in development^5,6,10,67^. However, current methods to evaluate drug response in PDCOs face several challenges: they can be destructive, overlook inter-PDCO heterogeneity, offer limited response metrics, or lack scalability. Common cell viability assays, such as tetrazolium and luciferase tests, assess metabolic function as a proxy for viable cell count but may introduce long-term toxicity, limit future use of precious samples, or fail to account for intra-tumoral heterogeneity^68–70^. Histological techniques yield detailed structural and molecular information but involve slow, destructive sample processing^71^. Traditional molecular biology methods (like RNA-seq, qPCR, and Western blots) capture treatment-induced changes in protein and RNA levels but often require sample pooling to obtain sufficient material^71^. While measuring PDCO diameter from bright-field images allows for monitoring the same PDCOs over time, this method provides only a single response metric (change in diameter) and can be subject to user bias due to manual execution^49,72^. Additionally, diameter changes are often preceded by more rapid metabolic responses^41^.

Performing optical redox imaging in a wide-field configuration is well suited for high-throughput applications in PDCO screening because of its simple, widely available instrumentation, large field-of-view, micron-scale resolution, and fast acquisition. Although WF ORI does not provide a readout at the cell-level, unlike two-photon ORI, our results indicate that the PDCO-level readout may better represent treatment response as WF ORI of individual PDCOs provided ∼7X increase in effect size compared to single cell two-photon ORI analysis for FOLFOX treatment of the same samples (**Fig. 2F**). This increase in effect size with WF ORI is likely due to the higher volume of cells interrogated with widefield illumination compared to a single z-slice with two-photon ORI. Two-photon z-stacks can be used to address this issue, with the caveat of lower throughput. WF ORI can screen >40 PDCOs / dish / 30s, providing greater power to resolve treatment effects compared to the lower-throughput two-photon ORI that screens 3-5 PDCOs / dish / 30min. Crucially for future dissemination of ORI measurements, WF ORI can be performed with hardware present in most research and clinical labs, at lower cost and higher throughput compared to two-photon ORI.

Furthermore, the isolation of ORR measurements to the leading-edge of the PDCOs, which corresponds to the proliferative cells with active metabolism, increases the sensitivity of ORR treatment response measurements (**Fig. 1C**). This is likely due to the removal of non-viable cells from the ORR measurement as most chemotherapies target rapidly dividing cells^22,62,63^. The components of FOLFOX, 5-FU and oxaliplatin^51,52^, both induce DNA damage, a common method for disrupting the growth of rapidly diving cancer cells. Similarly, panitumumab targets the EGF receptor^73^, which blocks a key pathway for cell growth. Though cells that are present in the core of the PDCOs will also be affected by treatment, the effect is diminished by the lack of rapid growth occurring in the internal PDCO niche as the cells are exposed to higher levels of metabolic waste (CO_2_ and lactate) and starvation due to lack of nutrient diffusion^62^. Additionally, the leading-edge ΔORR enhances differences in treatment response between responsive and resistant PDCOs within the same dish compared to more standard measurements of diameter change taken on the same PDCOs (**Fig. 4**). This resolution of single PDCOs is crucial as CRC samples are mutationally diverse and bulk sequencing does not capture rare alterations that often lead to tumor resistance^10,65^. Therefore, a tool that can quickly assess many PDCOs for driver mutations within days of PDCO culture (**Fig. 3**) and assess the efficacy of targeted therapies would be of great clinical use.

This study shows that wide-field optical redox imaging provides automated assessment of multiple quantitative variables at the PDCO level that capture treatment response and separate PDCO subpopulations based on key mutation phenotypes. An important output of this technical development is a framework to acquire robust redox images across multiple time points using a standard epifluorescence microscope. This proof-of-concept study used samples from a small subset of patients with known mutation profiles, and additional studies are needed to confirm that these specific observations apply to a variety of mutation profiles across additional patients. However, these methods can be adopted by non-experts to achieve reproducible images of treatment response in PDCOs using accessible instrumentation.

## 5. Conclusions

This study demonstrates a framework for performing wide-field optical redox imaging of PDCOs with the addition of leading-edge analysis that increases sensitivity to treatment response. Wide-field optical redox imaging is an easily scalable and accessible technology that provides functional metabolic information to characterize PDCO populations. Furthermore, this technique can detect differences in both baseline metabolism of PDCOs with altered driver mutations and differing responses to treatments with known mutational resistance. We believe that these approaches will enable broad dissemination of wide-field optical redox imaging to screen PDCOs, with applications in drug development and clinical treatment planning.

## 7. Data availability Statement

The data generated in this study are available upon request from the corresponding author.

## References

1. Fountzilas, E. & Tsimberidou, A. M. Overview of precision oncology trials: challenges and opportunities. Expert Rev. Clin. Pharmacol. 11, 797–804 (2018).

2. Prasad, V. Perspective: The precision-oncology illusion. Nature 537, S63–S63 (2016).

3. Tannock, I. F. & Hickman, J. A. Limits to Personalized Cancer Medicine. N. Engl. J. Med. 375, 1289–1294 (2016).

4. Letai, A. Functional precision cancer medicine—moving beyond pure genomics. Nat. Med. 23, 1028–1035 (2017).

5. Sachs, N. et al. A Living Biobank of Breast Cancer Organoids Captures Disease Heterogeneity. Cell 172, 373-386.e10 (2018).

6. Vlachogiannis, G. et al. Patient-derived organoids model treatment response of metastatic gastrointestinal cancers. Science 359, 920–926 (2018).

7. Saito, Y. et al. Establishment of Patient-Derived Organoids and Drug Screening for Biliary Tract Carcinoma. Cell Rep. 27, 1265-1276.e4 (2019).

8. Kopper, O. et al. An organoid platform for ovarian cancer captures intra- and interpatient heterogeneity. Nat. Med. 25, 838–849 (2019).

9. Sharick, J. T. et al. Cellular Metabolic Heterogeneity In Vivo Is Recapitulated in Tumor Organoids. Neoplasia 21, 615–626 (2019).

10. Sasaki, N. & Clevers, H. Studying cellular heterogeneity and drug sensitivity in colorectal cancer using organoid technology. Curr. Opin. Genet. Dev. 52, 117–122 (2018).

11. Benson, A. B. et al. Colon Cancer, Version 3.2024, NCCN Clinical Practice Guidelines in Oncology. J. Natl. Compr. Canc. Netw. 22, e240029 (2024).

12. Lubner, S. J., Uboha, N. V. & Deming, D. A. Primary and acquired resistance to biologic therapies in gastrointestinal cancers. J. Gastrointest. Oncol. 8, 499–512 (2017).

13. Pauli, C. et al. Personalized In Vitro and In Vivo Cancer Models to Guide Precision Medicine. Cancer Discov. 7, 462–477 (2017).

14. Ganesh, K. et al. A rectal cancer organoid platform to study individual responses to chemoradiation. Nat. Med. 25, 1607–1614 (2019).

15. Driehuis, E., Kretzschmar, K. & Clevers, H. Establishment of patient-derived cancer organoids for drug-screening applications. Nat. Protoc. 15, 3380–3409 (2020).

16. Ooft, S. N. et al. Patient-derived organoids can predict response to chemotherapy in metastatic colorectal cancer patients. Sci. Transl. Med. 11, eaay2574 (2019).

17. Tveit, K. M. & Pihl, A. Do cell lines in vitro reflect the properties of the tumours of origin? A study of lines derived from human melanoma xenografts. Br. J. Cancer 44, 775– 786 (1981).

18. Esquenet, M., Swinnen, J. V., Heyns, W. & Verhoeven, G. LNCaP prostatic adenocarcinoma cells derived from low and high passage numbers display divergent responses not only to androgens but also to retinoids. J. Steroid Biochem. Mol. Biol. 62, 391–399 (1997).

19. Daniel, V. C. et al. A Primary Xenograft Model of Small-Cell Lung Cancer Reveals Irreversible Changes in Gene Expression Imposed by Culture In vitro. Cancer Res. 69, 3364–3373 (2009).

20. Sambuy, Y. et al. The Caco-2 cell line as a model of the intestinal barrier: influence of cell and culture-related factors on Caco-2 cell functional characteristics. Cell Biol. Toxicol. 21, 1–26 (2005).

21. Wilding, J. L. & Bodmer, W. F. Cancer cell lines for drug discovery and development. Cancer Res. 74, 2377–2384 (2014).

22. Luengo, A., Gui, D. Y. & Vander Heiden, M. G. Targeting Metabolism for Cancer Therapy. Cell Chem. Biol. 24, 1161–1180 (2017).

23. Walsh, A. J. & Skala, M. C. Optical metabolic imaging quantifies heterogeneous cell populations. Biomed. Opt. Express 6, 559 (2015).

24. Varone, A. et al. Endogenous Two-Photon Fluorescence Imaging Elucidates Metabolic Changes Related to Enhanced Glycolysis and Glutamine Consumption in Precancerous Epithelial Tissues. Cancer Res. 74, 3067–3075 (2014).

25. Skala, M. C. et al. In vivo multiphoton microscopy of NADH and FAD redox states, fluorescence lifetimes, and cellular morphology in precancerous epithelia. Proc. Natl. Acad. Sci. 104, 19494–19499 (2007).

26. Shah, A. T., Diggins, K. E., Walsh, A. J., Irish, J. M. & Skala, M. C. In Vivo Autofluorescence Imaging of Tumor Heterogeneity in Response to Treatment. Neoplasia 17, 862–870 (2015).

27. Xu, H. N., Tchou, J., Feng, M., Zhao, H. & Li, L. Z. Optical redox imaging indices discriminate human breast cancer from normal tissues. J. Biomed. Opt. 21, 114003 (2016).

28. Alhallak, K. et al. Optical imaging of radiation-induced metabolic changes in radiation-sensitive and resistant cancer cells. J. Biomed. Opt. 22, 060502 (2017).

29. Ostrander, J. H. et al. Optical Redox Ratio Differentiates Breast Cancer Cell Lines Based on Estrogen Receptor Status. Cancer Res. 70, 4759–4766 (2010).

30. Kolenc, O. I. & Quinn, K. P. Evaluating Cell Metabolism Through Autofluorescence Imaging of NAD(P)H and FAD. Antioxid. Redox Signal. 30, 875–889 (2019).

31. Blacker, T. S. et al. Separating NADH and NADPH fluorescence in live cells and tissues using FLIM. Nat. Commun. 5, 3936 (2014).

32. Williamson, D. H., Lund, P. & Krebs, H. A. The redox state of free nicotinamide-adenine dinucleotide in the cytoplasm and mitochondria of rat liver. Biochem. J. 103, 514– 527 (1967).

33. Xiao, W., Wang, R.-S., Handy, D. E. & Loscalzo, J. NAD(H) and NADP(H) Redox Couples and Cellular Energy Metabolism. Antioxid. Redox Signal. 28, 251–272 (2018).

34. Quinn, K. P. et al. Quantitative metabolic imaging using endogenous fluorescence to detect stem cell differentiation. Sci. Rep. 3, 3432 (2013).

35. Georgakoudi, I. & Quinn, K. P. Optical Imaging Using Endogenous Contrast to Assess Metabolic State. Annu. Rev. Biomed. Eng. 14, 351–367 (2012).

36. Cannon, T. M., Shah, A. T., Walsh, A. J. & Skala, M. C. High-throughput measurements of the optical redox ratio using a commercial microplate reader. J. Biomed. Opt. 20, 010503 (2015).

37. Walsh, A. J. et al. Quantitative Optical Imaging of Primary Tumor Organoid Metabolism Predicts Drug Response in Breast Cancer. Cancer Res. 74, 5184–5194 (2014).

38. Shah, A. T., Heaster, T. M. & Skala, M. C. Metabolic Imaging of Head and Neck Cancer Organoids. PLOS ONE 12, e0170415 (2017).

39. Gillette, A. A. et al. Autofluorescence Imaging of Treatment Response in Neuroendocrine Tumor Organoids. Cancers 13, 1873 (2021).

40. Desa, D. E., Amitrano, M. J., Murphy, W. L. & Skala, M. C. Optical redox imaging to screen synthetic hydrogels for stem cell-derived cardiomyocyte differentiation and maturation. Biophotonics Discov. 1, (2024).

41. Gil, D. A., Deming, D. A. & Skala, M. C. Patient-derived cancer organoid tracking with wide-field one-photon redox imaging to assess treatment response. J. Biomed. Opt. 26, 036005 (2021).

42. Xu, H. N., Wu, B., Nioka, S., Chance, B. & Li, L. Z. Calibration of CCD-based redox imaging for biological tissues. in (eds. Hu, X.P. & Clough, A.V.) 72622F (Lake Buena Vista, FL, 2009). doi:10.1117/12.811769.

43. Ranji, M. et al. Fluorescence spectroscopy and imaging of myocardial apoptosis. J. Biomed. Opt. 11, 064036 (2006).

44. Schumacher, D. et al. Heterogeneous pathway activation and drug response modelled in colorectal-tumor-derived 3D cultures. PLOS Genet. 15, e1008076 (2019).

45. Phan, N. et al. A simple high-throughput approach identifies actionable drug sensitivities in patient-derived tumor organoids. Commun. Biol. 2, 78 (2019).

46. Favreau, P. F. et al. Label-free redox imaging of patient-derived organoids using selective plane illumination microscopy. Biomed. Opt. Express 11, 2591 (2020).

47. Xu, J. & Ying, W. Increased green autofluorescence is a marker for non-invasive prediction of H 2 O 2 -induced cell death and decreases in the intracellular ATP of HaCaT cells. Preprint at 10.1101/298075 (2018).

48. Surre, J. et al. Strong increase in the autofluorescence of cells signals struggle for survival. Sci. Rep. 8, 12088 (2018).

49. Pasch, C. A. et al. Patient-Derived Cancer Organoid Cultures to Predict Sensitivity to Chemotherapy and Radiation. Clin. Cancer Res. 25, 5376–5387 (2019).

50. Kratz, J. D. et al. Integrating Subclonal Response Heterogeneity to Define Cancer Organoid Therapeutic Sensitivity. Preprint at 10.1101/2021.10.15.464556 (2021).

51. Graham, M. A. et al. Clinical pharmacokinetics of oxaliplatin: a critical review. Clin. Cancer Res. Off. J. Am. Assoc. Cancer Res. 6, 1205–1218 (2000).

52. Joulia, J. M. et al. Plasma and salivary pharmacokinetics of 5-fluorouracil (5-FU) in patients with metastatic colorectal cancer receiving 5-FU bolus plus continuous infusion with high-dose folinic acid. Eur. J. Cancer Oxf. Engl. 1990 35, 296–301 (1999).

53. Stephenson, J. J. et al. An open-label clinical trial evaluating safety and pharmacokinetics of two dosing schedules of panitumumab in patients with solid tumors. Clin. Colorectal Cancer 8, 29–37 (2009).

54. Wagner, M. et al. Light Dose is a Limiting Factor to Maintain Cell Viability in Fluorescence Microscopy and Single Molecule Detection. Int. J. Mol. Sci. 11, 956–966 (2010).

55. Carpenter, A. E. et al. CellProfiler: image analysis software for identifying and quantifying cell phenotypes. Genome Biol. 7, R100 (2006).

56. Georgakoudi, I. & Quinn, K. P. Label-Free Optical Metabolic Imaging in Cells and Tissues. Annu. Rev. Biomed. Eng. (2023) doi:10.1146/annurev-bioeng-071516-044730.

57. Becker, W. The 10th TCSPC Handbook. vol. 10.

58. Walsh, A. & Skala, M. An automated image processing routine for segmentation of cell cytoplasms in high-resolution autofluorescence images. Prog. Biomed. Opt. Imaging -Proc. SPIE 8948, (2014).

59. Glass, G. V. Primary, Secondary, and Meta-Analysis of Research. Educ. Res. 5, 3 (1976).

60. Sawilowsky, S. S. New Effect Size Rules of Thumb. J. Mod. Appl. Stat. Methods 8, 597–599 (2009).

61. Dewhirst, M. W. & Secomb, T. W. Transport of drugs from blood vessels to tumour tissue. Nat. Rev. Cancer 17, 738–750 (2017).

62. Cui, X., Hartanto, Y. & Zhang, H. Advances in multicellular spheroids formation. J. R. Soc. Interface 14, 20160877 (2017).

63. Jagiella, N., Müller, B., Müller, M., Vignon-Clementel, I. E. & Drasdo, D. Inferring Growth Control Mechanisms in Growing Multi-cellular Spheroids of NSCLC Cells from Spatial-Temporal Image Data. PLoS Comput. Biol. 12, e1004412 (2016).

64. Walsh, A. J. et al. Optical Metabolic Imaging Identifies Glycolytic Levels, Subtypes, and Early-Treatment Response in Breast Cancer. Cancer Res. 73, 6164–6174 (2013).

65. Sforza, V. et al. Mechanisms of resistance to anti-epidermal growth factor receptor inhibitors in metastatic colorectal cancer. World J. Gastroenterol. 22, 6345 (2016).

66. DeStefanis, R. A. et al. Impact of baseline culture conditions of cancer organoids when determining therapeutic response and tumor heterogeneity. Sci. Rep. 12, 5205 (2022).

67. Tiriac, H. et al. Organoid Profiling Identifies Common Responders to Chemotherapy in Pancreatic Cancer. Cancer Discov. 8, 1112–1129 (2018).

68. Hamid, R., Rotshteyn, Y., Rabadi, L., Parikh, R. & Bullock, P. Comparison of alamar blue and MTT assays for high through-put screening. Toxicol. In Vitro 18, 703–710 (2004).

69. Fotakis, G. & Timbrell, J. A. In vitro cytotoxicity assays: Comparison of LDH, neutral red, MTT and protein assay in hepatoma cell lines following exposure to cadmium chloride. Toxicol. Lett. 160, 171–177 (2006).

70. Van Tonder, A., Joubert, A. M. & Cromarty, A. D. Limitations of the 3-(4,5-dimethylthiazol-2-yl)-2,5-diphenyl-2H-tetrazolium bromide (MTT) assay when compared to three commonly used cell enumeration assays. BMC Res. Notes 8, 47 (2015).

71. Lee, S. H. et al. Tumor Evolution and Drug Response in Patient-Derived Organoid Models of Bladder Cancer. Cell 173, 515-528.e17 (2018).

72. Fricke, S. L. et al. MTORC1/2 Inhibition as a Therapeutic Strategy for PIK3CA Mutant Cancers. Mol. Cancer Ther. 18, 346–355 (2019).

73. Addeo, R. et al. Panitumumab: a new frontier of target therapy for the treatment of metastatic colorectal cancer. Expert Rev. Anticancer Ther. 10, 499–505 (2010).

