## supplemental table for "Wide-field optical redox imaging with leading-edge detection for assessment of patient-derived cancer organoids"

**Supplemental Video 1.** Method of spiking individual PDCO's for mutation mixing studies

**Supplemental Table 1.** Number of PDCO's assessed with WF ORI per experiment

| Experiment | PDCO Mutations | # Patient Samples | Control | FOLFOX | Panitumumab |
|---|---|---|---|---|---|
| FOLFOX | Wild-type | 2 | 175 | 205 | |
| Mutation Profiling | WildType | 4 | 128 | | |
| | KRAS + / PIK3CA - | 3 | 199 | | |
| | KRAS - / PIK3CA + | 2 | 226 | | |
| | KRAS + / PIK3CA + (DM) | 2 | 493 | | |
| Targeted Treatment | WildType | 1 | | | 27 |
| | KRAS + / PIK3CA + (DM) | 1 | | | 24 |

**Supplemental Table 2.** Number of PDCOs and cells assessed with two-photon ORI per condition

| Experiment | PDCO Mutations | # Patient Samples | Sample Level | Control | FOLFOX |
|---|---|---|---|---|---|
| FOLFOX | Wild- type | 2 | PDCOs | 8 | 8 |
| | | | Cells | 1974 | 1644 |
